# Class-imbalanced Unsupervised and Semi-Supervised Domain Adaptation for Histopathology Images

**DOI:** 10.1101/2023.03.08.531734

**Authors:** S. Maryam Hosseini, Abubakr Shafique, Morteza Babaie, H.R. Tizhoosh

**Affiliations:** Kimia Lab, University of Waterloo, ON, Canada; Vector Institute, MaRS Centre, Toronto, Canada; Rhazes Lab, the Department of Artificial Intelligence and Informatics, Mayo Clinic, Rochester, MN, USA

**Keywords:** Deep Learning, Histopathology, Whole Slide Images, Unsupervised Domain Adaptation, Semi-supervised Domain Adaptation, Class imbalance

## Abstract

In dealing with the lack of sufficient annotated data and in contrast to supervised learning, unsupervised, self-supervised, and semi-supervised domain adaptation methods are promising approaches, enabling us to transfer knowledge from rich labeled source domains to different (but related) unlabeled target domains, reducing distribution discrepancy between the source and target domains. However, most existing domain adaptation methods do not consider the imbalanced nature of the real-world data, affecting their performance in practice. We propose to overcome this limitation by proposing a novel domain adaptation approach that includes two modifications to the existing models. Firstly, we leverage the focal loss function in response to class-imbalanced labeled data in the source domain. Secondly, we introduce a novel co-training approach to involve pseudo-labeled target data points in the training process. Experiments show that the proposed model can be effective in transferring knowledge from source to target domain. As an example, we use the classification of prostate cancer images into low-cancerous and high-cancerous regions.

## I. Introduction

The remarkable recent success of deep learning has shown great promise in digital pathology which arises from the co-operations of deep learning (DL) experts and histopathologists, leading to advanced ML methods for whole slide image (WSI) analysis. Despite the success of DL models in medical applications, their prediction performance degrades if the characteristics of training and testing data distribution diverge. This divergence is most probably inevitable in histopathology image analysis because tissue preparation protocols, staining procedures, and digital scanners vary between pathology labs. Additionally, even in the same pathology laboratory, these processes are likely to evolve in time by changing scanners or tissue preparation protocols. Therefore, WSIs, as the digital images in pathology, exhibit considerable variations between labs and also within labs, that can be called “*domain shift*”. As a result, any DL model trained on a labeled dataset from one domain may not have acceptable performance on data from another domain. This apparently happens because extracted deep features from convolution neural networks have transitioned from general to domain-specific, especially in higher layers [1].

One naive solution to the domain shift challenge is to annotate the data of the new domain and fine-tune the DL model, i.e., the deep network. However, data annotation is an expensive, time- and resource-consuming process. To alleviate the need for larger annotated datasets, unsupervised domain adaptation (UDA) and semi-supervised domain adaptation (SDA) are emerging as potential remedies. The intuition of domain adaptation is to involve domain adaptation methods in the process of learning the representations and deep features.

Domain adaptation [2] is a promising scheme allowing us to train the model on labeled domain (called *source domain*) while performing well on different (but related) unlabeled (or limited labeled) domain (called *target domain*). Domain adaptation is supposed to mimic the human vision system, which can transfer learned knowledge from one or multiple source domains to another related (but different) domain. Similar to the human vision system, this method enables the model to learn generalized image representations that are task-specific and domain invariant.

Many real-world datasets are naturally imbalanced with respect to class-instance distribution. Some classes are under-represented and rare, while others are dominant and contain many data points. This makes DL models biased toward the dominant class, preventing them from learning minor classes. The class-imbalance issue makes the domain adaptation methods more challenging since the approaches need to deal with data distribution divergence between domains while considering the class imbalance nature of the source and target data.

In this paper, we address the class-imbalance challenge in domain adaptation employing two alterations to existing models. We use the focal loss for the label classification error estimation to train the model.

## II. Background

Generally, deep networks rely on abundant labeled training instances, while both train and test datasets are assumed to have the same distributions. However, collecting large labeled dataset is a time- and resource-consuming process. Additionally, it is likely that train and test data have domain shift. Transfer learning is a promising method to solve these challenges, helping to transfer knowledge across domains. The main goal of transfer learning is to improve the prediction performance on target domains by transferring knowledge from different but related source domains [3].

The first subcategory is “unsupervised” transfer learning, which refers to process that labelled information is unavailable in both source and target domains. The second subcategory is “transductive” transfer learning, where label information is not available or is limited at target, while it is available at source domain [4]. In this subcategory, considering only one single task, it is called *domain adaptation*. In domain adaptation, one or more labeled source domains may be adopted, transferring knowledge to unlabeled or limited labeled target domain. In the third subcategory, the label information is available at both source and target domains.

Considering label sets at source and target domain, there are four categories of domain adaptation problems, closed set, open set, partial set, and universal set domain adaptation [5]. In the closed set domain adaptation, both source and target domains share the same labels. In the open set domain adaptation, the source label set is a subset of the target label set [6]. Partial set domain adaptation is the opposite of open set domain adaptation, meaning that the target label set is a subset of the source label set [7]. Finally, universal set domain adaptation generalizes the above scenarios, requiring no knowledge about source and target label sets relation. In this scenario, source and target data may share common label sets, and also each domain may have a particular label set or outlier classes [5].

In this paper, we consider the “closed set domain adaptation” problem, where both source and target data points share the same label set.

Domain adaptation can be accomplished in either one step or multi-steps. In one-step domain adaptation, it is assumed that both source and target domains are directly related, such that knowledge can be transferred in one step. However, this assumption does not hold all the time. In some scenarios, source and target domains are not directly related; thus, one step knowledge transfer may not be effective. It requires some intermediate domains to bring source and target domains together. In this paper, one-step domain adaptation method has been considered since images from different hospitals are directly related.

### A. Unsupervised Domain Adaptation

The unsupervised domain adaptation (UDA) has emerged as a promising learning approach to address the lack of annotated data, becoming prominent in transferring knowledge between domains. There are three lines of research on UDA methods [1], [8], [9]: 1) statistical-based, 2) reconstruction-based, and 3) adversarial-based UDA. The common idea of all these methods is to involve domain adaptation in the training process, such that the extracted features are domain invariant but still class discriminative.

#### 1) Statistical-based UDA

The statistical-based domain adaptation methods rely on statistics to decrease the distribution difference between source and target data. These methods involve fine-tuning the deep network using target data points. The common methods in statistical-based include maximum mean discrepancy (MMD), correlation alignment (CORAL), batch normalization (BN), entropy minimization (EM), moment matching, and Wasserstein discrepancy [9].

- **MMD** [10] is a kernel method, estimating sample differences. Relying on kernel function, MMD is a metric to compare the distributions between two datasets. In neural networks, MMD is estimated between the representation of each domain, reducing their differences in the feature space.
- **CORAL** is another statistical method to domain adaptation. The main goal of CORAL is to learn linear transforms to align the second order statistics (covariances) of source and target domains [11].
- **BN** [12] aims to decrease the covariance shift across domains using BN layers. There are previous works that leverage BN layers, called DM-layer, specifically for domain adaptation.
- **EM** is a common objective loss function in domain adaptation. It estimates the distance between probability distributions of domains, encouraging low entropy and consistency on domain predictions [13].
- **moment matching** [14], [15] focuses on the moments of extracted features. Moment matching estimates one or more moments of features, estimating the discrepancy of feature distributions across domains to lessen the inconsistency between domains.
- **Wasserstein metric** is another statistical method in domain adaptation. It is a metric to measure discrepancy across domains, finding domain invariant and class discriminative representations.

#### 2) Reconstruction-based UDA

The reconstruction-based domain adaptation attempts to diminish the gaps between domains by reconstructing all data samples, mapping both source and target data into a shared feature space. The reconstruction-based method preserve distinctiveness of intra-domain representations while focuses on the indistinguishably of inter-domain representations. The reconstruction-based methods include encoder-decoder models, dictionary and sparse coding models. [9].

- In **encoder-decoder models**, one idea is to adapt the encoder based on data points from one domain and learn decoder to reconstruct data points in another domain [16]. In this approach, the encoder learns to map images to domain-invariant features, while decoder learns domain-specific features to reconstruct images from domain-invariant features of the encoder. Another idea is to use stacked auto-encoders, which uses encoder for representation learning and decoder for reconstruction [17]. There are also other works that leverage auto-encoder to learn self-domain and between domain reconstruction.
- **Dictionary and sparse coding models** involve updating and adapting a dictionary to represents data of both source and target domains [18].

#### 3) Adversarial-based UDA

This line of research is among the recent deep learning methods for domain adaptation. These methods leverage adversarial objective to reduce the distribution difference across domains via a domain discriminator. The domain discriminator is a binary classifier, determining whether a data point belongs to source or target domain. The binary domain discriminator encourages domain confusion using an adversarial objective function. The adversarial methods can be classified into two main subcategories, generative and non-generative [1].

- **Generative models** have been inspired by generative adversarial networks (GAN) [19]. These methods employ GANs, learning to generate target samples from source data points or noise vectors.
- **Non-generative models** aim to learn domain-invariant representations. These methods leverage domain confusion loss to learn features. The non-generative adversarial domain adaptation was proposed [2] by adding a binary domain discriminator module to reduce the gap between domains in feature space.

### B) Semi-supervised Domain Adaptation

Semi-supervised learning is a method somewhere between supervised (that rely on labeled data for training) and unsupervised methods (that rely on unlabeled data for training). Semi-supervised domain adaptation can be accomplished by involving labeled target data points in each of the previously mentioned UDA approaches. Each of the previously mentioned methods are applicable in semi-supervised learning by involving the labeled target data in training. Despite many studies in domain adaptation, there are a few research studies on domain adaptation with imbalance datasets [20]. Most methods assume class balanced samples.

### C) Class Imbalanced Dataset

The class imbalance is a quite common issue in almost all real-world datasets. The imbalance makes the DL models biased toward the dominant class, ignoring minority (small/rare) classes. In medical imaging, the minority class may be of great importance since, for instance, some rare cancer subtypes may be misclassified if the model is incapable of dealing with the imbalance. Therefore, having such a bias increases the danger of miscategorizing minor yet critical samples of rare classes. There are various methods to combat the disproportionate number of class instances. These methods can be classified into three categories [21], data-level methods, algorithm-level methods, and hybrid methods.

Data-level methods include up-sampling and down-sampling. Up-sampling is a procedure to expand the minority class by building synthetic data points. Injecting these synthetic data points into the pool of data helps to create a more balanced dataset. Unlike up-sampling that focuses on the minority class, down-sampling is concerned with the majority class. Down-sampling selects a subset of the majority samples for training the model to avoid any bias. Neither of these methods is favorable. Down-sampling discards samples of the dominant class, losing some available information. Up-sampling generates samples similar to the minor class, resulting in over-fitting [22], [23].

The algorithm-level methods leave the dataset untouched and focus on changing the learning algorithm to control the unbalanced nature of the dataset, being unbiased toward any major class. The algorithm-level methods include modifying the learning by altering the loss function to address the class imbalance dataset. These loss functions enable samples of rare classes to contribute more to the loss compared to the data points that belong to the majority classes [24], [25].

The hybrid methods are combinations of data-level and algorithm-level methods. These methods slightly touch training data samples to create a more balanced dataset, as well as modify learning procedures to have unbiased training [24]. In this paper, we employ focal loss in our training to estimate classification error [25] as well as pseudo-labeling samples to improve training.

## III. Method

The overview of the proposed method has been depicted in Figure 1. In this section, each part of this framework will be explained.

**Fig. 1:**
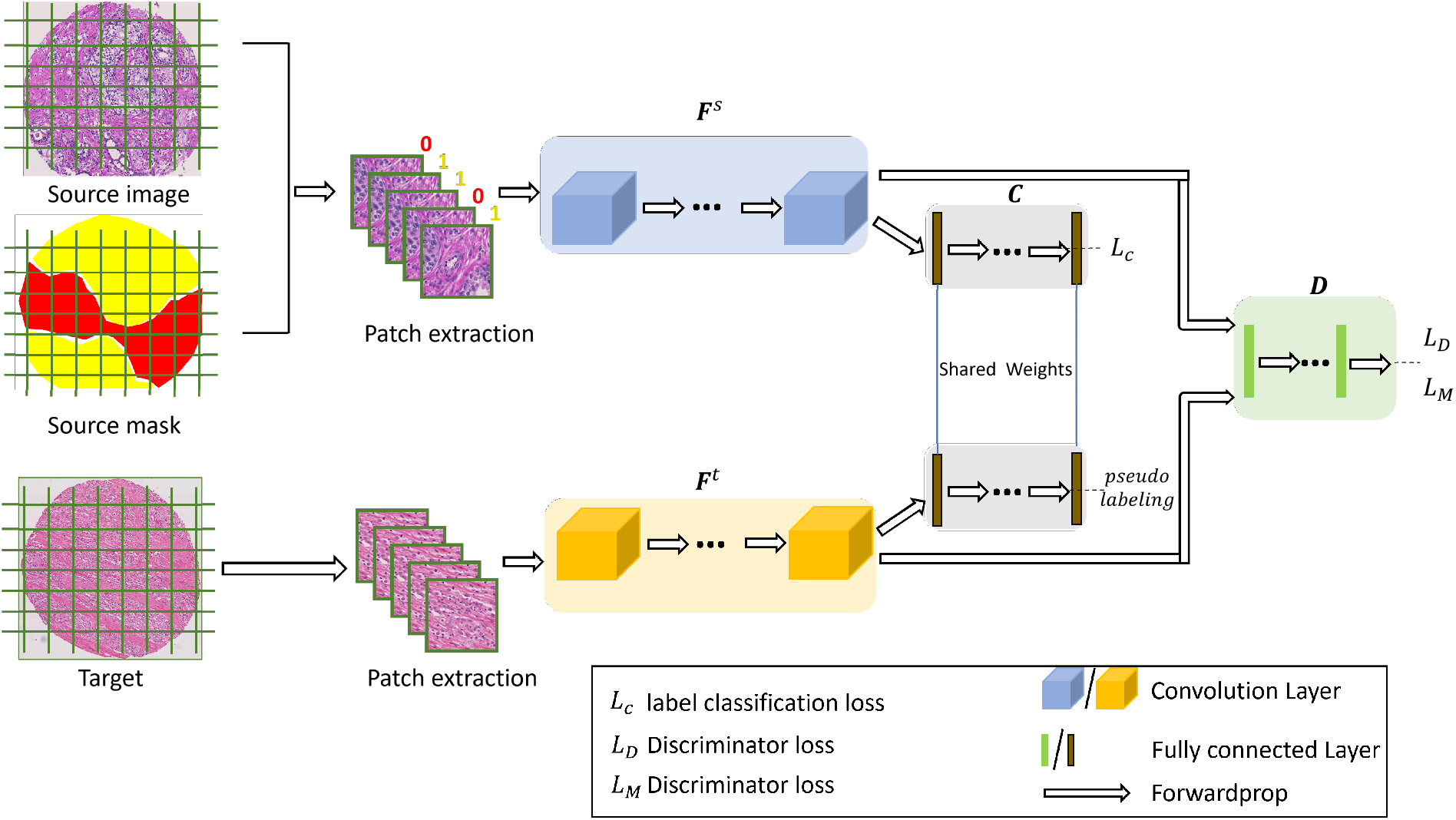
Training model using focal loss and pseudo-labels. Overview of the proposed methodology. The features of the source and target images are extracted using **F**^*s*^ and **F**^*t*^ networks, respectively. Using labeled samples of the source domain and pseudo-labeled samples of the target domain, we estimate classification loss for each domain employing the classification network **C**. Additionally, the domain classifier network **D** provides us with domain discriminator loss estimations.

### 1) Problem formulation

In this section, necessary notations and definitions will be introduced. The source domain includes *N*_*s*_ labeled samples denoted by 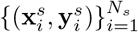 where 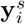 is a one-hot vector, representing the label of the *i*th sample of the source domain. The target domain includes *N*_*t*_ unlabeled data samples denoted by 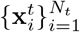 and 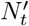 labeled samples denoted by 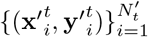 where 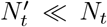. We leverage only *N*_*t*_ unlabeled target samples for UDA, and both *N*_*t*_ unlabeled and 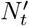 labeled samples for SDA. As illustrated in Figure 1, we extract features of source and target samples using two separate networks, **F**^*s*^ with parameters *θ*^*s*^ and **F**^*t*^ with parameters *θ*^*t*^. The label classifier **C** with parameters *θ*^*c*^ allows us to perform classification, estimating label classification loss. The domain binary classifier **D** with parameters *θ*^*d*^ enables us to estimate domain discriminator loss, helping us to train the network in an adversarial manner.

### 2) Co-Training

To improve training, unlabeled samples of the target domain will be involved in training to contribute to label classification loss, called co-training. In co-training, labels are assigned to the target data points, called *pseudo-labels*, selecting the most reliable pseudo-labeled target data points that meet three criteria to become part of training samples in the next iteration. Given target data point 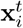, its pseudo-label is defined as

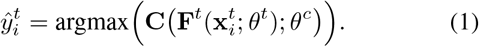

The first criterion to select a pseudo-labeled target sample 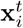 is to have a high classification score. The classification score is defined as the maximum prediction probability

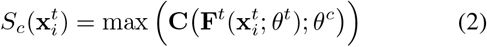

To have reliable co-training data, only pseudo-labeled samples with high prediction probability will be selected. Therefore, the first necessary condition to select a pseudo-labeled sample 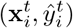 is 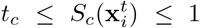, where the constant *t*_*c*_ is the minimum acceptable prediction probability.

The second criterion to select a pseudo-labeled sample 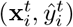 is domain discriminator score 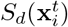, which is the probability that the sample 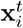 belongs to the source domain. This metric is defined as

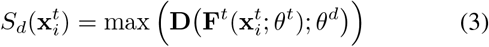

whereas sample 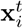 is a suitable candidate if the domain discriminator is confused and have probability near 0.5 for the domain of 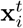. Therefore, the second necessary condition is 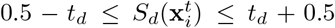, where the constant *t*_*d*_ determines the margin from 0.5.

The third criterion to select a pseudo-labeled sample 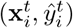 is the imbalance score. Using this metric, samples that belong to rare minor classes will be chosen. For a given data point 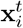 with pseudo-label 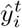, the imbalance score is defined as

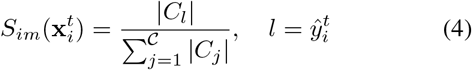

where *C* is the total number of classes, |*C*_*j*_| indicates the size of the *j*th class, and index *l* represents the class to which 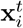 belong. Obviously, small 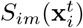 indicates that 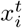 belongs to a rare class. Therefore, the third necessary condition is 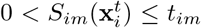 whereas the constant *t*_*im*_ enables us to select those pseudo-labeled target data points that belongs to minor classes. The combination of labeled source data points and selected pseudo-labeled target data points is denoted by 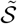. In summary, a pseudo-labeled target data point 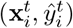 will be selected for training if all the three constraints beloware satisfied:

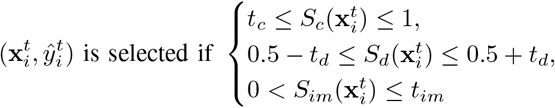

### 3) Loss Function

Using labeled source samples and selected pseudo-labeled target data points, label classification loss ℒ_*c*_ is estimated by employing focal loss rather than cross-entropy loss. Focal loss is the modified version of the standard cross-entropy loss, first proposed in [25] to boost the detection of rare samples. It reduces the loss associated with well-classified samples while increasing the loss of minor class data points. This loss function prevents dominant class samples from overwhelming the classifier during training, boosting unbiased training. In this proposal, the focal loss [25] will be leveraged for classification error estimation to combat the class imbalance challenge. Figure 2 represents the difference between cross-entropy loss and focal loss. As can be seen, the focal loss function is a special form of cross-entropy loss. Unlike cross-entropy loss function, samples have different contributions in the focal loss. The contribution of each sample depends on its prediction probability. The contribution of the well-classified samples to ultimate loss is less than other samples. As shown in Figure 2, the derivative of the focal loss function is less than the cross-entropy loss function for well-classified samples. However, the derivative of the focal loss is more than cross-entropy when prediction probability is low and samples are hard to classify. Mathematically, the focal loss is defined as [25]

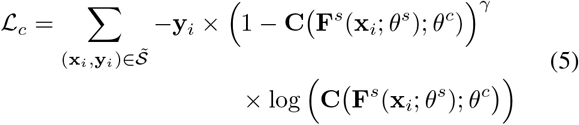

where the constant *γ* is focusing parameter that adjusts the down-weighted rate of samples with low prediction probability. As shown in [25], *γ* ∈ [0.5, 5] provides us with more robust result. In our experiments, we consider *γ* = 2.

**Fig. 2:**
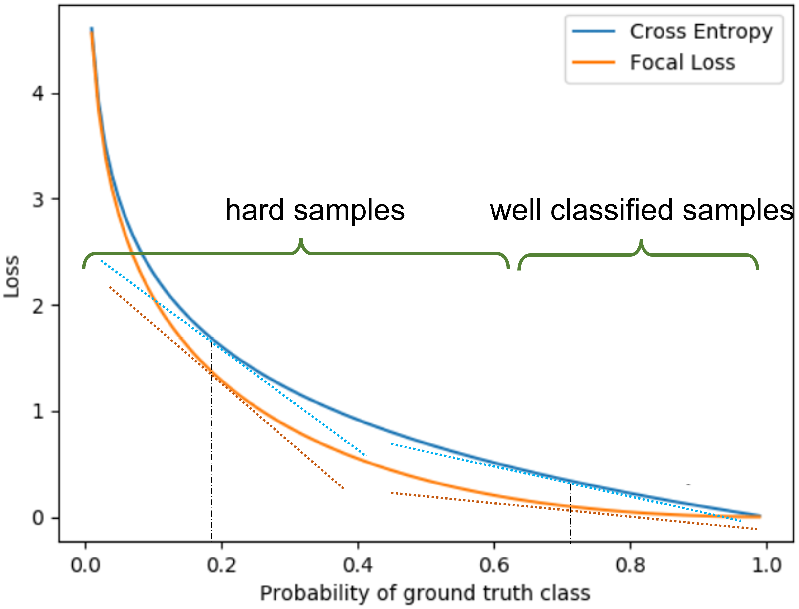
Focal loss function v.s. cross entropy loss function.

The adversarial domain adaptation is performed by using domain discriminator **D**. The main goal of the using domain discriminator is to mix data points of both domains such that they they seem to belong to one domain. The adversarial domain loss is defined as [2]

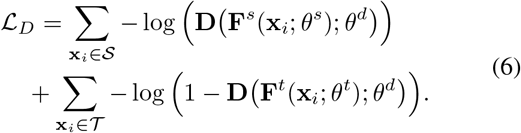

Another adversarial loss function that has been used is [2]

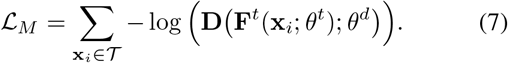

## IV. Experiments and Results

In this section, we present our experimental results.

### 1) Dataset

We consider two pairs of datasets to evaluate the proposed approach for transferring knowledge from one domain to another for the prostate cancer classification task.

The first pair consists of two tissue micro-array (TMA) datasets [26], [27]. One sample image from each of these datasets has been shown in Fig. 3. The dataset [26] has been prepared and published by Harvard university. This dataset consists of 886 tissue micro-arrays of prostate cancer. TMAs are 3100 × 3100 images at 40x magnification with Hematoxylin and Eosin (H&E) staining. TMAs have pixel-level annotations corresponding to Gleason Scores provided by expert pathologists. For the simplicity, this dataset is denoted by TMA2 in the rest of this paper. The dataset [27] has been published in MICCAI 2019 as a “Grand Challenge” for pathology. This dataset consists of 244 TMA images at 40x magnification with H&E staining. The size of the most TMA images is 5120 × 5120, but some images are slightly smaller or larger. The TMA images are annotated by expert pathologists. The annotation is in form of pixel-level Gleason Score label and core-level Gleason Score label. In the evaluation, only pixel-level annotations have been used. For simplicity, this dataset is denoted by TMA1 in the rest of this paper.

**Fig. 3:**
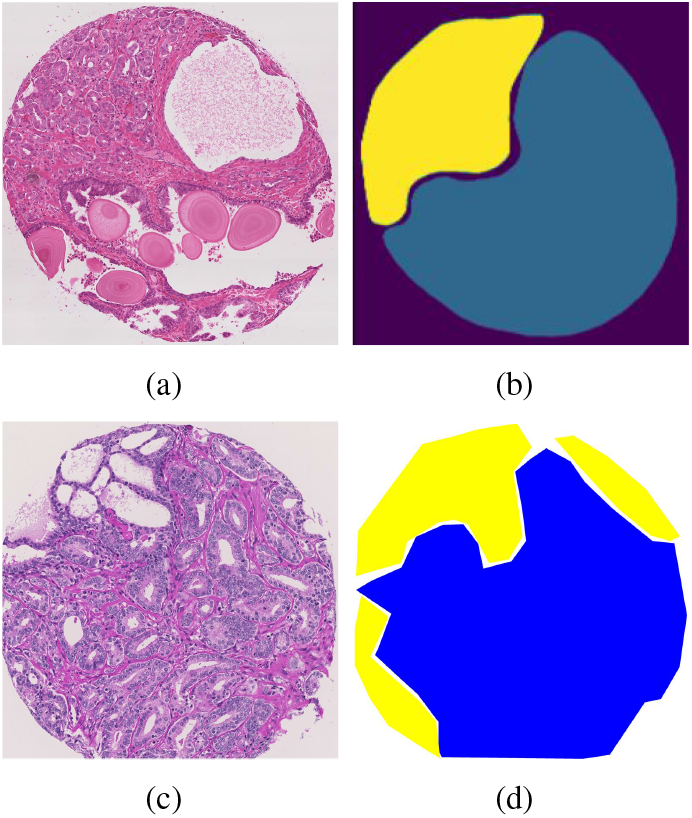
TMA Datasets Visualization. Samples with their masks from tissue micro-array datasets, TMA1 and TMA2 datasets. Images in the first and second row belong to TMA1 and TMA2 respectively.

**Fig. 4:**
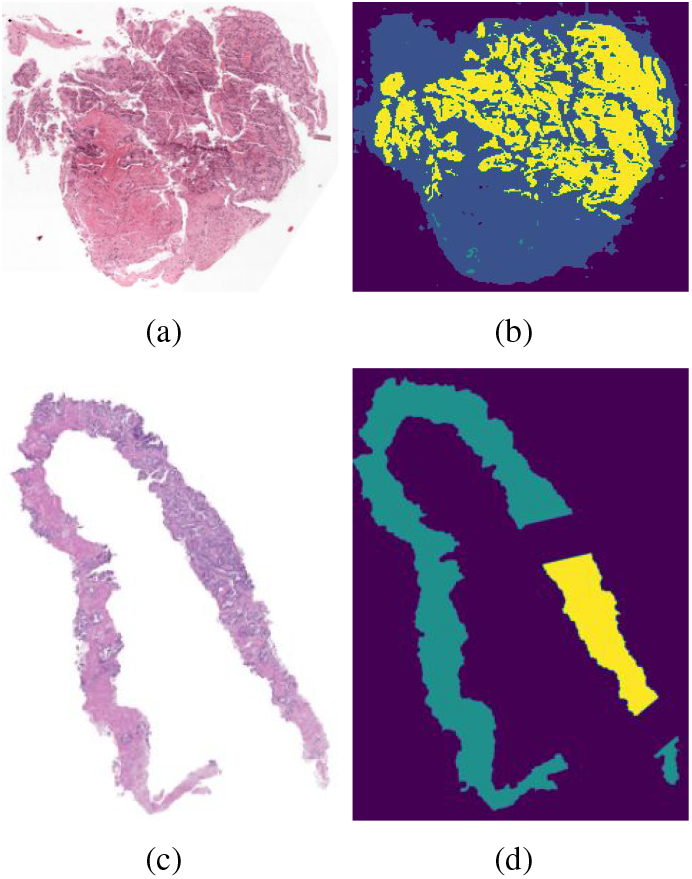
WSI Datasets Visualization. Samples with their masks from PANDA dataset. Images in PANDA dataset are provided by two organizations, Radboud and Karolinska. Images in the first row belong to Radboud organization while images in the second row provided by Karolinska.

TMA images of both datasets have been cropped to patches of 740 × 740 pixels for training and test purposes. Table I represents the details of each datasets. A label (Gleason Score) has been assigned to each of 740 × 740 patches based on the majority of the pixel annotation. Patches with background or benign label have been discarded. The Gleason Score 3 is considered as less cancerous, while Gleason Score 4 and 5 are regarded as more cancerous.

**TABLE I:** The brief overview of the source and target datasets used in our experiments. (IF: Imbalance Factor)

| Dataset | IF | # Img | # Patches | Tail/Head size |
| --- | --- | --- | --- | --- |
| PANDA/ Karolinska | 2.48 | 5414 | 10460 | 7460/ 3000 |
| PANDA/ Radboud | 8.07 | 5160 | 21698 | 2391/ 19308 |
| TMA1/ Gleason2019 | 2.98 | 244 | 4546 | 1142/ 3404 |
| TMA2/ Harvard | 1.16 | 641 | 5620 | 2594/ 3026 |

For the second dataset pair, the validation is performed on the Prostate cANcer graDe Assessment (PANDA) dataset [28], introduced in MICCAI 2020 challenge. PANDA is the largest available dataset of prostate cancer annotated with the ISUP Grade that represents the severity of prostate cancer. In this dataset, labels range from 0 to 5, where 0 represents non-cancer cases, 1 and 5 belong to the lowest and the most severe cancer cases, respectively. There are also noisy per-pixel segmentation masks depicting image regions led to the ISUP Grade. This dataset includes H&E stained whole slide images of prostate tissue, provided by two pathology labs, namely Radboud University Medical Center and Karolinska Institute. The dataset from Karolinska Institute has been considered as source domain with labeled data and data from Radboud University Medical Center as target domain without the label. The WSIs have been cropped in patches of size 512 × 512 for training and testing purposes. The patches with either tissue area less than the half of the image size or no segmentation masks have been eliminated. A label has been assigned to each patch according to the majority vote of the pixel level mask annotation. The ISUP Grades 0, 1, 2 are considered as low cancerous category, while ISUP Grades 3, 4 and 5 as high cancerous group.

According to [29], imbalance factor (IF) of a dataset has been defined, representing to waht degree a data set is imbalanced. The IF of a given data set is the ratio of the size of the dominant class to the size of the smallest class, which specifies the maximum between-class imbalance level expressed as 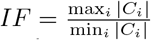. Clearly, the larger IF, the more imbalanced the dataset.

The four publicly available datasets with different IF values have been used to examine the performance of proposed approach for knowledge transfer when there is class imbalance. Table III provides a brief review of these datasets. The fist two datasets consist of WSIs of prostate cancers whereas the last two datasets consist of TMA images of prostate cancers.

### 2) Implementation

The ultimate goal is to train the model that could classify WSIs into low and high ISUP Grade/Gleason Scores at target domain. Therefore, WSIs have been divided into two groups at both source and target domains, considering WSIs with ISUP Grades (Gleason Scores) 0, 1, 2 (score 3) as low grade while WSI with ISUP Grades (Gleason Scores) 3, 4, 5 (scores 4, and 5) as high grade. The patches have been resized to 256 × 256 pixels and fed into feature extractor networks. DenseNet121 [30] followed by batch normalization and convolution layers have been used as the feature extractor, **F**^**s**^ and **F**^**t**^. The label classifier **C** choose to have three fully connected layers, estimating label classification error. The binary domain classifier **D** choose to have two fully connected layers, estimating domain discriminator loss. The parameters of the pseudo-labeling have been chosen as *tc* = 0.95, *t*_*d*_ = 0.4, *t*_*i*_ = 0.4. In each iteration, target samples that meet these criteria will be selected and added to source training samples in the next training iteration. The network has been trained in 20 epochs, using a learning rate of 0.0001.

The performance of the proposed framework has been compared with baseline and two other methods [2], [31]. The “baseline” is a simple DenseNet121 that has been fine-tuned on only source data samples. The performance has been evaluated using three performance metrics, majority-minority different (M-MD), mean per class accuracy (MCA), and accuracy. The M-MD and MCA performance metrics are specifically used for imbalanced datasets.

The experimental results of the proposed method are presented in Table II. As can be seen, the proposed UDA has promising performance compared to the baseline and two other reference methods. Additionally, the proposed SDA method with 10% and 20% labeled target data samples outperforms the other methods. We have not reported the experimental results of transferring knowledge from Karolinska to Radboud since the annotated samples of Karolinska seem to be noisy. Therefore, the model cannot learn transferable knowledge from this dataset.

**TABLE II:** Comparison of classification using three metrics, the majority-minority difference (M-MD), mean per class accuracy (MCA), and accuracy (Acc) are in percent.

| TMA2 → TMA1 | | | |
| --- | --- | --- | --- |
| Algorithm | M-MD ↓ | MCA ↑ | Acc ↑ |
| Baseline | 21.34 | 71.56 | 66.85 |
| Ganin et al. [2] | 5.36 | 72.02 | 69.43 |
| Ren et al. [31] | 9.12 | 70.3 | 66.82 |
| **Ours (UDA)** | 4.67 | 72.54 | 72.89 |
| **Ours (SDA, 10%)** | 4.34 | 73.72 | 73.39 |
| **Ours (SDA, 20%)** | 4.12 | 73.94 | 74.87 |
| TMA1 → TMA2 | | | |
| Algorithm | M-MD ↓ | MCA ↑ | Acc ↑ |
| Baseline | 44.65 | 65.61 | 67.28 |
| Ganin et al. [2] | 42.34 | 70.83 | 70.31 |
| Ren et al. [31] | 43.67 | 65.93 | 68.72 |
| **Ours (UDA)** | 20.34 | 73.67 | 74.86 |
| **Ours (SDA, 10%)** | 19.45 | 76.87 | 75.46 |
| **Ours (SDA, 20%)** | 18.91 | 77.12 | 75.92 |
| Radboud → Karolinska | | | |
| Algorithm | M-MD ↓ | MCA ↑ | Acc ↑ |
| Baseline | 60.31 | 65.34 | 70.49 |
| Ganin et al. [2] | 58.54 | 70.45 | 71.57 |
| Ren et al. [31] | 55.39 | 70.13 | 72.41 |
| **Ours (UDA)** | 18.45 | 73.24 | 74.68 |
| **Ours (SDA, 10%)** | 16.94 | 76.78 | 76.12 |
| **Ours (SDA, 20%)** | 16.05 | 77.65 | 76.94 |

### 3) Ablation study

We conducted an ablation study. The results of this study have been reported in Table III that shows co-training improves the classification performance.

**TABLE III:** Ablation study. Accuracy comparison for **TMA1** → **TMA2**. All numbers are in percent.

|  | Co-training<br>with CE loss | Only Focal loss | Co-training<br>with Focal loss |
| --- | --- | --- | --- |
| Accuracy ↑ | 72.77 | 70.55 | 74.86 |
| M-MD ↓ | 21.53 | 34.56 | 20.34 |
| MCA ↑ | 72.26 | 70.96 | 73.67 |

## V. Conclusions

One of the main challenges of using deep learning in digital pathology is the scarcity of annotated histopathology datasets. To overcome this shortage, we proposed a domain adaptation method for the classification of WSI into low-cancerous and high-cancerous regions. Relying on the domain adaptation, we want to transfer knowledge from the label-rich source domain to the unannotated target domain. We considered the realistic practical assumption that both source and target datasets are class imbalanced. We adopted *focal loss* in the training rather than the cross-entropy loss, focusing attention on samples belong to rare classes and allowing samples from minor classes to contribute more to the loss. We also proposed a novel *co-training* approach to enrich the training data. In this approach, we assign labels to unlabeled target data points, selecting the most reliable ones for training. The validation results demonstrated competitive performance on public datasets.

